# Juxtacap Nucleotide Sequence Modulates eIF4E Binding and Translation

**DOI:** 10.1101/165142

**Authors:** Heather R. Keys, David M. Sabatini

## Abstract

mRNA translation is an energetically costly activity required for almost all biological processes. The multiprotein complex eIF4F, which bridges the 5’ cap and the polyA tail through eIF4E and eIF4G, respectively, is necessary for efficient translation initiation of most mRNAs and is an important target of translational control. Previous work suggests that cap-proximal nucleotides can modulate eIF4E binding to mRNAs, but the effect of specific cap-proximal nucleotide sequences on eIF4E recruitment and the ultimate consequences for translation remain unknown. Using RNA Bind-n-Seq on a model 5’ UTR, we systematically identify eIF4E-intrinsic cap-proximal nucleotide binding preferences. mRNAs with highly-bound motifs are translated well in a cell-free system, whereas those with low-eIF4E-binding motifs are not. However, eIF4E juxtacap motif preferences do not dictate the ribosome occupancy of endogenous mRNAs in cells, suggesting that the effect of juxtacap sequence on eIF4E binding and translation is mRNA-context-dependent. Accordingly, a single downstream point mutation that disrupts a predicted base pair with a preferred juxtacap nucleotide increases translation. The juxtacap sequence is a previously unappreciated determinant of eIF4E recruitment to mRNAs, and we propose that differences in mRNA 5’ end accessibility defined by the juxtacap sequence are important for establishing translational efficiency.

## INTRODUCTION

Protein synthesis is a fundamental process essential for almost all biological phenomena and as such is highly regulated. As the rate-limiting step of protein synthesis under normal conditions (1,2), translation initiation can be dynamically regulated in a global and general manner through the modulation of core translation factors (reviewed in (3,4)). Additionally, the translation of specific mRNAs can be temporally and spatially controlled in diverse cell types or at specific developmental stages (reviewed in (3,4)).

Two main regulatory arms impact translation initiation in eukaryotes. One point of regulation occurs at the formation of the ternary complex, which comprises the initiation factor eIF2, the initiator methionyl-tRNA, and GTP, and which is required for recognition of the start codon (reviewed in (4)). The other major point of regulation is at the recruitment of the 43S preinitiation complex (43S PIC) to the mRNA (reviewed in (3)). The 43S PIC includes the 40S small ribosomal subunit, the ternary complex, and initiation factors eIF1, eIF1A, and eIF3 (reviewed in (3,5)). The 43S PIC is recruited to the mRNA via an interaction between eIF3 and the multiprotein complex eIF4F (6). The mechanistic target of rapamycin complex 1 (mTORC1), a kinase complex that integrates nutrient and growth factor inputs to promote several processes important for cell growth, regulates the assembly of eIF4F to impact translation initiation (reviewed in (7)).

eIF4F comprises the cap-binding protein eIF4E, the scaffolding protein eIF4G, and the helicase eIF4A (reviewed in (8)). eIF4E performs the key first step of eIF4F function by recognizing the 5’ end of the mRNA via its interaction with the 7-methylguanosine (^me7^G) cap, thereby positioning eIF4F and the 43S PIC at the 5’ end of the mRNA (9). Incorporation of eIF4E into the eIF4F complex is an important modulatory point for translational regulation, and is the step specifically controlled by mTORC1 (10,11). mTORC1 phosphorylates the 4E-binding proteins (4EBPs), rendering them inactive for binding to eIF4E (10). Because the 4EBPs compete with eIF4G for binding to eIF4E (11), mTORC1 activity promotes assembly of eIF4F. eIF4G is a large protein that binds eIF4E, mRNA, polyA-binding protein (PABP), eIF3, and eIF4A (reviewed in (12)). eIF4G performs several functions: stabilizing the RNA-eIF4E interaction by binding RNA directly, bridging the 5’ and 3’ termini of the mRNA through interactions with eIF4E and PABP, and recruiting the 43S PIC to the 5’ end of the mRNA via the interaction with eIF3 (reviewed in (12)). Furthermore, eIF4G localizes eIF4A to the mRNA to unwind 5’UTR secondary structure and facilitate scanning of the 40S ribosomal subunit (reviewed in (12)). Thus, eIF4F coordinates many interactions required for translation initiation.

eIF4E is the least abundant member of eIF4F (13,14) and, upon mTORC1 inhibition, becomes even more limited due to sequestration from eIF4F by the 4EBPs (10,11,15). eIF4E is overexpressed in several types of cancer (reviewed in (16)), experimental overexpression of eIF4E in cells leads to oncogenic transformation *in vitro* (17), and overexpression in mice leads to increased tumor incidence (18). Intriguingly, eIF4E overexpression affects translation of some mRNAs more than others (19), suggesting that translation of this subset of messages is particularly dependent on eIF4E. One class of mRNAs whose translation is enhanced by eIF4E overexpression harbors a pyrimidine stretch at the 5’ terminus (5’ TOPs) (19). mRNAs harboring this 5’ TOP motif are also the mRNAs whose translation is the most strongly inhibited when eIF4F is disrupted by acute inhibition of mTOR (20,21). The sensitivity of 5’ TOP mRNA translation to eIF4F inhibition and eIF4E overexpression suggests that the nucleotides proximal to the 5’ cap may play a role in the recruitment of eIF4E to mRNAs.

Biochemical and structural work has primarily detailed the eIF4E-cap interaction (22-27), and the role of cap-proximal nucleotides in eIF4E recruitment remains unclear. The crystal structure of eIF4E bound to the cap analog ^me7^GDP reveals a cleft of positive electrostatic potential directly adjacent to ^me7^GDP, through which the +1 nucleotide could be threaded (22). In the co-crystal structure of human eIF4E with the cap analog ^me7^GpppA, contact of eIF4E with adenosine at the +1 position stabilizes an otherwise flexible C-terminal loop of eIF4E, indicating that eIF4E can contact at least the first juxtacap nucleotide (25). NMR experiments with yeast eIF4E and ^me7^GpppA support this finding (23). However, in a co-crystal structure of murine eIF4E with ^me7^GpppG, no electron density is observed for the +1 guanosine, indicating the absence of a direct interaction with the +1 nucleotide (26), and suggesting that an interaction between eIF4E and the +1 nucleotide could depend on nucleotide identity.

How eIF4E binding and translation are influenced by additional nucleotides downstream of the +1 position is even less clear. Subsequent cap-proximal nucleotides can decrease the binding of eIF4E relative to the 5’ cap alone, if those nucleotides induce secondary structure in the RNA near the 5’ end that physically blocks eIF4E binding (28). In another case, cap-proximal secondary structure does not prevent eIF4E binding, but may change the trajectory of the RNA as it exits the cap-binding pocket of eIF4E in a manner inhibitory for translation (29). Conversely, kinetic measurements have shown that the presence of a short oligonucleotide stretch adjacent to the cap can increase the affinity of eIF4E for the RNA (30). It appears that the presence of juxtacap nucleotides can influence the binding of eIF4E to mRNA; however, the direction, magnitude, and sequence-dependence of this effect remain unclear.

Here, we systematically identify juxtacap sequences that affect recruitment of eIF4E to a model 5’ UTR, and that modulate translation of mRNAs containing the model 5’UTR in a cell-free system. We propose that the juxtacap sequence influences eIF4E binding and translation indirectly, by influencing secondary structure at the 5’ end of the mRNA. Mutations downstream of the juxtacap motif predicted to alter the cap-proximal secondary structure affect translation *in vitro*, in the absence of changes to the juxtacap motif itself. The juxtacap sequence is a previously unappreciated determinant of eIF4E recruitment to mRNAs, and could be an important contributor to establishing translational efficiency *in vivo*.

## MATERIALS AND METHODS

### Reagents

Enzymes were purchased from New England Biolabs, except where noted. Primers were obtained from Integrated DNA Technologies. Antibiotics and chemicals were purchased from Sigma, except where noted. TBE-Urea gels, TBE gels, and sample loading dyes were purchased from ThermoFisher. DMEM and inactivated fetal bovine serum (IFS) were purchased from US Biologicals.

### eIF4E cloning, expression, and purification

Murine eIF4E was amplified by PCR from a plasmid encoding eIF4E (Addgene #38239; mouse and rat eIF4E sequences are identical), cloned into the pET302/NT-His bacterial expression vector (ThermoFisher) downstream of a 6xHis tag using XhoI and BamHI restriction sites, and clones were sequence verified using the T7 Promoter Primer.

meIF4E_Nterm_F:AATTCGCTCGAGatggcgactgtcgaaccg

heIF4E_Nterm_R:GCAGCCGGATCCttaaacaacaaacctatt

T7 Promoter Primer: TAATACGACTCACTATAGGG

A streptavidin-binding peptide (SBP) tag was introduced in place of the 6xHis tag using a geneblock (IDT) encoding SBP with a 5’ NdeI site and a 3’ XhoI site.

SBP_geneblock_NdeI_XhoI:

GATATACATATGGACGAAAAAACGACCGGGTGGCGTGGCGGCCATGTCGTGGAGGGCCTGGCAGGCGAGCTGGAACAACTGCGCGCACGTCTGGAACACCATCCTCAGGGACAGCGCGAACCAGTGAATTCGCTCGAGatggcg

Sequence-verified clones were transformed into Rosetta (DE3) pLysS cells (EMD Millipore), inoculated into 50 ml LB supplemented with 34 μg/ml chloramphenicol and 100 μg/ml ampicillin, and grown overnight at 37°C. Cultures were diluted 1:100 in LB with chloramphenicol and ampicillin and grown at 37C until the OD was between 0.5 and 0.7. Cultures were shifted to 16°C for 20 minutes and eIF4E expression was induced for 18 hours at 16°C by addition of 0.5 mM IPTG (FisherScientific). All subsequent purification steps were performed at 4°C. Cells were lysed in Resuspension Buffer (20 mM HEPES pH 7.4, 100 mM KCl, 2 mM EDTA, 1 mM DTT) three times in a French press, and CHAPS was immediately added to 0.3%. Lysates were sonicated twice and were cleared sequentially by low speed centrifugation and ultracentrifugation.

^me7^GTP beads (preparation described below) were washed once with Wash Buffer (20 mM HEPES pH 7.4, 100 mM KCl, 0.3% CHAPS, 1 mM DTT) and added to the cleared lysate. Binding was allowed to proceed for 90 minutes at 4°C with gentle mixing. Beads were washed 3 times for 5 minutes each with 20 ml Wash Buffer and SBP-eIF4E was eluted with three consecutive washes for 15 minutes each with 1 ml Elution Buffer (20 mM HEPES pH 7.4, 500 mM KCl, 0.3% CHAPS, 1 mM DTT). Eluates were pooled and desalted using a PD-Miditrap G-25 size exclusion column (GE Healthcare) into Wash Buffer. Protein was concentrated in an Amicon Ultra centrifugal filter (EMD Millipore) with a molecular weight cutoff of 10,000 kDa. Protein was centrifuged at 17,000xg for 10 minutes at 4°C, and the soluble material was stored at 4°C and used within 3 days. Protein was determined to be pure by Coomassie staining. Protein was centrifuged again for 10 minutes at 17,000xg at 4C immediately prior to use, and only the soluble material was used.

### Preparation of ^me7^GTP beads

^me7^GTP beads were prepared as described previously (31), with slight modifications. Briefly, one molar equivalent of sodium meta-periodate (FisherScientific) in 0.1 M NaOAc pH 6 was added to ^me7^GTP in water and incubated for 90 minutes at 4°C. Adipic acid dihydrazide agarose beads (Sigma) were washed once with water and with 0.1 M NaOAc pH 6. Beads were resuspended in 0.1 M NaOAc pH 6 and transferred to the tube containing the oxidized ^me7^GTP. The mixture was incubated with rotation for 90 minutes at 4°C, and sodium cyanoborohydride was added. The beads were incubated overnight at 4C with mixing, and washed several times with 1M NaCl and Storage Buffer (50 mM HEPES pH 7.4, 100 mM KCl, 0.2 mM EDTA). Beads were stored at 4°C in 10 ml storage buffer with 0.02% sodium azide.

### Rps20 5’UTR cloning

The murine Rps20 5’UTR was cloned from mouse embryonic fibroblast cDNA into pIS1-Eef2-5UTR-renilla (Addgene plasmid #38235) by PCR amplification with the following Rps20 5’UTR-specific primers containing a 5’ EcoRI restriction site and modified T7 RNA Polymerase promoter sequence (containing no guanosines at positions +1, +2, and +3), and a 3’ NheI restriction site.

mRps20_dbTOP_0G_F: ttggcgaattc<u>TAATACGACTCACTATA</u>cTTTCTGAGCCCCGGCGG mRps20_R: tggtggctagcGGCGCGGCTTCCTGACCG

### *In vitro* transcription, RNA processing, and purification

The DNA template for the randomized Bind-n-Seq library was generated by PCR amplification of the Rps20 5’UTR from pIS1-mRps20_dbTOP_0G using an Rps20 5’UTR-specific forward primer containing a 5’ EcoRI restriction site, a modified T7 RNA Polymerase promoter containing only a +1 guanosine, randomized nucleotides at positions +2 through +6, and an Rps20 5’UTR-specific reverse primer containing the endogenous murine Rps20 Kozak sequence.

mRps20_db_G1N9_5U_F:ttggcgaattcTAATACGACTCACTATAGNNNNNNNNNcccggcggtgcgcg mRps20_Kozak_R: GGCTGAggcgcggcttcctgaccg

An aliquot of the PCR product was run on an agarose gel to confirm the amplification of a single product prior to purification of the rest of the PCR product using the Quiaquick gel extraction kit (Qiagen). PCR products were eluted in DEPC-treated water, and all subsequent processing steps were carried out using RNAse-free reagents.

The Bind-n-Seq RNA library was transcribed from the DNA template at 37C for two hours in a reaction containing 1X NEB T7 RNA Polymerase buffer, 100 mM DTT, 0.6 U SuperaseIn (ThermoFisher), 2 mM ATP, 2 mM UTP, 2 mM CTP, 2 mM GTP, 1 ug randomized DNA template, and 2.5 U T7 RNA Polymerase per 100 μl reaction volume. DNAse was added directly to the reaction at a concentration of 10 U/100 μl and the reactions were incubated for an additional 30 minutes at 37°C. The transcription reactions were extracted with acid phenol/chloroform (FisherScientific), and samples were incubated 5 minutes at room temperature with periodic vortexing. The aqueous layer was extracted with chloroform and the RNA was precipitated three times with NH_4_OAc and isopropanol.

The RNA was denatured at 75°C for 10 minutes and put on ice, then capped *in vitro* for two hours at 37°C using the ScriptCap m^7^G Capping System (Cellscript) according to manufacturer’s instructions. Capping reactions were extracted with acid phenol/chloroform and precipitated three times with NH_4_OAc and isopropanol.

RNA was treated sequentially with 5’ Polyphosphatase (Epicentre) and Terminator 5’ Phosphate-Dependent Exonuclease (Epicentre) to remove uncapped RNA. 5’ Polyphosphatase reactions contained 1X Polyphosphatase Buffer and 0.6 U 5’ Polyphosphatase per 1 μg RNA (approximately 20 U 5’ Polyphosphatase per 1 nmol RNA). RNA was denatured for 5 minutes at 75°C and placed on ice just prior to addition to the reaction. Reactions were incubated at 37°C for 2 hours, and the RNA precipitated with NaOAc and isopropanol. 5’ Polyphosphatase-treated RNA was denatured for 5 minutes at 75°C and digested with Terminator Exonuclease in a reaction containing 1X Terminator Reaction Buffer A and 0.3U Terminator Exonuclease per 1 ug RNA for 2 hours at 30°C. EDTA was added to 5mM to stop the reaction, and RNA was precipitated with NaOAc and isopropanol. RNA was resuspended in DEPC water, TBE-Urea sample buffer was added, and samples were denatured 10 minutes at 75°C and placed on ice. Samples were loaded onto a prerun 6% TBE-Urea gel and run for 50 minutes at 200V. Gels were stained 5 minutes with SYBR gold (FisherScientific) at 1:12,500 in TBE, and gel slices were crushed by centrifugation at 4°C at 17,000xg for 3 minutes through a 0.65 ml microfuge tube pierced with an 18G needle. RNA was eluted in 300 mM NaOAc, 1 mM EDTA overnight at 4°C with agitation. Eluate was filtered through a SpinX 0.22 μM filter (Sigma), and RNA was precipitated with isopropanol. RNA was resuspended in DEPC water, and the RNA concentration was measured by absorbance at 260 nm.

### RNA Bind-n-Seq

RNA-Bind-n-Seq was adapted from the method of Lambert *et al*. (32). Each 250μl binding reaction contained 1X Binding Buffer (20 mM HEPES pH 7.4, 150 mM KCl, 2.5 mM MgCl_2_, 0.3% CHAPS, 0.5 mM DTT), SBP-eIF4E (at a concentration of 50 nM, 125 nM, 250 nM, 750 nM, 1 μM, or 1.25 μM), randomized RNA library at a fixed concentration of 9 μM, 15 ng/μl polyI RNA, and 0.4 U/μl SuperaseIn. Control reactions also contained either 100 μM GTP or 100 μM^me7^ GTP. SBP-eIF4E was equilibrated in the reaction mix without RNA library for 30 minutes at 23°C, after which the RNA library was added and the binding reaction allowed to proceed for one hour at 23°C with constant mixing at 750 rpm in an Eppendorf Thermomixer. During the binding reaction, 100 μl streptavidin magnetic beads per reaction were washed in a batch 3 times with 1 ml 1X Binding Buffer. Beads were resuspended in 1X Binding Buffer and aliquotted to one tube per reaction. The remaining buffer was removed just prior to addition of each binding reaction; the binding reaction and bead mixture was incubated for an additional one hour at 23°C with constant mixing at 750 rpm. Reactions were washed once with 1 ml 1X Wash Buffer (20 mM HEPES pH 7.4, 150 mM KCl, 0.5 mM EDTA, 0.3% CHAPS) and resuspended in 250 μl Elution Buffer (20 mM HEPES pH 7.4, 1 mM EDTA, 1% SDS). Samples were heated for 10 minutes at 70°C, and eluate was transferred to a new tube. Samples were extracted with acid phenol/chloroform at 65°C for 5 minutes with periodic vortexing. Samples were put on ice for 5 minutes prior to recovery of the aqueous layer. The aqueous layer was extracted with acid phenol/chloroform a second time for 5 minutes at room temperature with periodic vortexing. The aqueous layer was then extracted with one volume chloroform for 30 seconds with constant vortexing, and the RNA was precipitated with NaOAc and isopropanol and resuspended in 20 μl 10 mM Tris pH 8 per sample.

### Sequencing Library Preparation

Sequencing library preparation was performed similarly to Ingolia *et al*. (33). cDNA synthesis was performed with Superscript III Reverse Transcriptase (ThermoFisher). The reverse transcription primer contained a 5’ phosphate to allow enzymatic circularization of the resulting single-stranded DNA, and also a stretch of 6 random nucleotides, to ensure complexity in base composition in the first several cycles during sequencing. The RT primer also contained the reverse complement of the 5’ Illumina adapter, an abasic site to allow relinearization of the circularized single-stranded DNA to increase PCR efficiency if necessary, and the reverse complement of the 3’ Illumina adapter. Finally, the RT primer contained the reverse complement of the 3’ end of the Rps20 5’UTR, which is shared by all RNAs in the library.

dbRps20_RT primer:

/5Phos/NNNNNNGATCGTCGGACTGTAGAACTCTGAAC/iSp18/CACTCA/iSp18/ccttggcacccgagaattcca GGCTGAggcgcggcttcctg

For the binding reaction samples, 10 μl RNA was used in a 20 μl cDNA synthesis reaction; for the input library, 0.5 pmol RNA in 10 μl 10 mM Tris pH 8 was used. The RNA was mixed with 1 μl 50 μM RT primer and 1 μl 10 mM dNTP mix. Reactions were incubated for 5 minutes at 65°C and placed on ice. To each chilled reaction, a mix containing 0.5 μl DEPC water, 4 μl 5X First Strand Synthesis Buffer, 1 μl 0.1 M DTT, 0.5 μl SuperaseIn, and 1 μl Superscript III Reverse Transcriptase was added. Reactions were incubated for 45 minutes at 48°C. RNA was hydrolyzed by addition of 2.2 μl 1 M NaOH and incubation for 20 minutes at 98°C, followed by addition of 2.2 μl 1 M HCl. DNA was recovered by precipitation with NaCl and isopropanol and resuspended in 10 μl 10 mM Tris pH 8. DNA was loaded onto a pre-run 6% TBE-Urea gel and run for 25 minutes at 200 V. Gels were stained with SYBR Gold, and gel slices were incubated in 300 mM NaCl with 1 mM EDTA and 10 mM Tris pH 8 overnight at 4°C. Eluate was filtered through a SpinX 0.22 μM filter and precipitated with isopropanol.

Pellets were resuspended in 4.5 μl Circularization mix without CircLigase (1X CircLigase Buffer, 50 μM ATP, 2.5 mM MnCl_2_, and 50 U CircLigase; Epicentre). CircLigase was added, and reactions were incubated for one hour at 60°C. CircLigase was heat inactivated by incubation for 10 minutes at 80°C. DNA was precipitated with NaCl and isopropanol, and resuspended in 12 μl 10 mM Tris pH 8.

The randomized sequences and a portion of the Rps20 5’UTR were PCR amplified using a forward primer containing the 5’ Illumina adapter sequence, and one of several Illumina Tru-Seq small RNA reverse primers containing the Illumina 3’ adapter sequence and a unique barcode.

randomized_PCR_F: AATGATACGGCGACCACCGA<u>GATCTACAC</u>**GTTCAGAGTTCTACAGTCCGA**

Illumina Index Primer 1 (1.25μM eIF4E + GTP): CAAGCAGAAGACGGCATACGAGAT**CGTGAT**GTGACTGGAGTT**CCTTGGCACCCGAGAATTCCA**

Illumina Index Primer 2 (1.25μM eIF4E + ^me7^GTP): CAAGCAGAAGACGGCATACGAGAT**ACATCG**GTGACTGGAGTT**CCTTGGCACCCGAGAATTCCA**

Illumina Index Primer 3 (50nM eIF4E): CAAGCAGAAGACGGCATACGAGAT**GCCTAA**GTGACTGGAGTT**CCTTGGCACCCGAGAATTCCA**

Illumina Index Primer 4 (125nM eIF4E): CAAGCAGAAGACGGCATACGAGAT**TGGTCA**GTGACTGGAGTT**CCTTGGCACCCGAGAATTCCA**

Illumina Index Primer 5 (250nM eIF4E): CAAGCAGAAGACGGCATACGAGAT**CACTGT**GTGACTGGAGTT**CCTTGGCACCCGAGAATTCCA**

Illumina Index Primer 7 (750nM eIF4E): CAAGCAGAAGACGGCATACGAGAT**GATCTG**GTGACTGGAGTT**CCTTGGCACCCGAGAATTCCA**

Illumina Index Primer 9 (1μM eIF4E): CAAGCAGAAGACGGCATACGAGAT**CTGATC**GTGACTGGAGTT**CCTTGGCACCCGAGAATTCCA**

Illumina Index Primer 10 (1.25μM eIF4E): CAAGCAGAAGACGGCATACGAGAT**AAGCTA**GTGACTGGAGTT**CCTTGGCACCCGAGAATTCCA**

Illumina Index Primer 11 (input library): CAAGCAGAAGACGGCATACGAGAT**GTAGCC**GTGACTGGAGTT**CCTTGGCACCCGAGAATTCCA**

For each sample, five reactions of 16.7 μl each containing 1X Phusion HF Buffer, 200 μM of each dNTP, 500 nM each of forward and reverse primer, 1 μl circularized ssDNA, and 0.334 U Phusion High-Fidelity DNA Polymerase were heated for 30 seconds at 98°C and amplified for 6, 8, 10, 12, or 14 cycles with the following program:

1. 98°C 10 seconds
2. 60°C 10 seconds
3. 72°C 10 seconds

Reactions were run on a pre-run 8% TBE gel for one hour at 180 V. DNA was extracted from bands where the amplification had not reached saturation and was extracted from the gel slices and precipitated as described above. Libraries were resuspended in 5 μl water per slice and were pooled and sequenced on an Illumina HiSeq Sequencer according to standard procedures.

### Data Analysis

Motifs were extracted from raw sequencing data using a custom Python script to identify reads containing the sequence “GAGCCCCG,” which is shared by all 5’UTRs in the library and is directly downstream of the random 5-mer. Reads that did not contain the constant sequence or that did not contain an entire 5-mer preceding the constant sequence were discarded. The five-nucleotide motifs directly preceding the constant sequence were extracted and their frequencies calculated for each binding reaction. All subsequent data manipulation and statistical analysis was performed in R.

### Cell-Free Translation Assay

The cell-free translation assay was adapted from Rakotondrafara *et al*. (34). Briefly, 10x10^6^ Hela cells were seeded in 15 cm dishes in 20 ml DMEM supplemented with 10% IFS and penicillin/streptomycin. The following day, cells were washed with PBS, trypsinized, and resuspended in ice cold DMEM without serum. Cells were pelleted for 3 min at 318xg at 4°C. Cells were washed once with 10 ml ice cold PBS and pelleted again. Residual PBS was removed, and cells were resuspended in one pellet volume Hypotonic Lysis Buffer (16 mM HEPES pH 7.4, 10 mM KOAc, 0.5 mM MgOAc, 5 mM DTT) with one freshly added EDTA-free protease inhibitor tablet (Roche) per 10 ml buffer. Cells were incubated on ice for 10 minutes and homogenized by passage through a 27G1/2 needle 15 times at 4°C. Lysate was cleared by centrifugation for 1 minute at 13,300xg at 4°C, and soluble material was normalized to 10 mg/ml, aliquotted, and stored at -80°C.

mRNAs encoding the Rps20 5’UTR with various juxtacap motifs were generated by PCR amplification of pIS1-Rps20_dbTOP_0G using unique forward primers to introduce different juxtacap motifs, together with the mRps20_R primer.

mRps20_GGCGT: ttggcgaattcTAATACGACTCACTATAg**GGCGT**GAGCCCCGGCGG

mRps20_GGTGC: ttggcgaattcTAATACGACTCACTATAg**GGTGC**GAGCCCCGGCGG

mRps20_GGCGA: ttggcgaattcTAATACGACTCACTATAg**GGCGA**GAGCCCCGGCGG

mRps20_GCGTT: ttggcgaattcTAATACGACTCACTATAg**GCGTT**GAGCCCCGGCGG

mRps20_GGCGC: ttggcgaattcTAATACGACTCACTATAg**GGCGC**GAGCCCCGGCGG

mRps20_GCGTA: ttggcgaattcTAATACGACTCACTATAg**GCGTA**GAGCCCCGGCGG

mRps20_GCTTT: ttggcgaattcTAATACGACTCACTATAg**GCTTT**GAGCCCCGGCGG

mRps20_GGCTT: ttggcgaattcTAATACGACTCACTATAg**GGCTT**GAGCCCCGGCGG

mRps20_AAGCC: ttggcgaattcTAATACGACTCACTATAg**AAGCC**GAGCCCCGGCGG

mRps20_AGGCT: ttggcgaattcTAATACGACTCACTATAg**AGGCT**GAGCCCCGGCGG

mRps20_ATGCC: ttggcgaattcTAATACGACTCACTATAg**ATGCC**GAGCCCCGGCGG

mRps20_AGGCC: ttggcgaattcTAATACGACTCACTATAg**AGGCC**GAGCCCCGGCGG

mRps20_TGGCT: ttggcgaattcTAATACGACTCACTATAg**TGGCT**GAGCCCCGGCGG

mRps20_AGGTC: ttggcgaattcTAATACGACTCACTATAg**AGGTC**GAGCCCCGGCGG

mRps20_AGACC: ttggcgaattcTAATACGACTCACTATAg**AGACC**GAGCCCCGGCGG

mRps20_AAGCT: ttggcgaattcTAATACGACTCACTATAg**AAGCT**GAGCCCCGGCGG

mRps20_TTGCT: ttggcgaattcTAATACGACTCACTATAg**TTGCT**GAGCCCCGGCGG

mRps20_TTGCC: ttggcgaattcTAATACGACTCACTATAg**TTGCC**GAGCCCCGGCGG

The PCR products were digested with NheI and EcoRI and ligated into the pIS1-Rps20_dbTOP_0G vector upstream of the *Renilla* luciferase gene and an encoded polyA tail. Sequence-verified clones were linearized by BamHI digest and transcribed *in vitro* as described above for the Bind-n-Seq library, except that mRNAs were co-transcriptionally capped using anti-reverse cap analog (ARCA; New England Biolabs), instead of post-transcriptionally capped. After extraction with acid phenol/chloroform and precipitation with NaOAc and isopropanol, mRNAs were gel separated on a pre-run 6% TBE-Urea gel for 2.5 h at 200 V. mRNAs were extracted from the gel slices as described above, and resuspended in 10 mM Tris pH 8. mRNAs were stored at -80C, and working stocks were stored at -20°C.

10 ng of each mRNA was translated in a 10 μl reaction containing 40 mM KOAc, 2 mM MgCl_2_, 0.8 mM ATP, 0.1 mM GTP, and 1X translation buffer (16 mM HEPES pH 7.4, 20 mM creatine phosphate, 0.1 μg/μl creatine kinase, 0.1 mM spermidine (freshly diluted from 1 mM stock), and amino acids at RPMI concentrations) for 30 minutes at 37°C. Translation was detected using the *Renilla* Luciferase Assay System (Promega). 10 μl 1X *Renilla* Luciferase Lysis Buffer was added on ice to stop the reactions. 10μl of each reaction was added to 50 μl of a *Renilla* Luciferase Assay Buffer and *Renilla* Luciferase Assay Substrate mixture immediately prior to reading for 10 seconds in a luminometer. Luciferase readings for experimental samples were background-subtracted using readings from samples containing 10 mM Tris pH 8 instead of RNA.

### Statistical Analyses and Data Visualization

The following R packages were used for data visualization and statistical analyses (R Core Team (2015) R: A language and environment for statistical computing. R Foundation for Statistical Computing, Vienna, Austria. https://www.R-project.org/):

seqLogo (Bembom,O. (2016) seqLogo: Sequence logos for DNA sequence alignments. R package version 1.34.0.)

colorspace (Ihaka,R., Murrell,P., Hornik,K., Fisher,J.C. and Zeileis,A. (2015). colorspace: Color Space Manipulation. R package version 1.2-6. http://CRAN.R-project.org/package=colorspace)

plot3D (Soetaert,K. (2014) plot3D: Plotting multi-dimensional data. R package version 1.0-2. http://CRAN.R-project.org/package=plot3D)

scales (Wickham,H. (2015) scales: Scale Functions for Visualization. R package version 0.3.0. http://CRAN.R-project.org/package=scales)

flux (Jurasinski,G., Koebsch,F., Guenther,A. and Beetz,S. (2014) flux: Flux rate calculation from dynamic closed chamber measurements. R package version 0.3-0. http://CRAN.R-project.org/package=flux)

### Transcriptional Start Site and Ribosome Footprinting Analysis

The FANTOM5 project used cap analysis of gene expression (CAGE) to identify robust transcriptional start sites (TSSs) from numerous mouse and human samples (35). TSS data from NIH3T3 cells (36) was processed to identify dominant TSSs using the CAGEr package (37) (with default parameters, except fitRange was set from 10 to 1000 and slope was set to 1.14) and subsetted by TSSs beginning with a guanosine using custom R scripts. TSSs beginning with a guanosine were mapped to overlapping genes using the BEDTools suite (38). Ribosome occupancy of these genes was extracted from ribosome footprinting data from p53-/-mouse embryonic fibroblasts (21). eIF4E binding was plotted against ribosome occupancy for each juxtacap motif represented in a TSS that overlaps with a gene present in the ribosome footprinting dataset.

### RNA Secondary Structure Prediction

RNA secondary structures of model 5’UTRs were predicted using RNAfold (39), and base pairing probabilities were extracted for nucleotides at the 5’ end of the RNA. A nucleotide at a particular position was deemed inaccessible if it had a base pairing probability greater than 0.75.

## RESULTS

### eIF4E RNA Bind-n-Seq

To systematically identify juxtacap sequences that affect eIF4E binding, we performed a modified RNA Bind-n-Seq (RBNS) protocol (32) using a library of 7-methylguanosine-capped (^me7^G-capped) RNAs adapted from the murine Rps20 5’ UTR (Figure 1A). The T7 RNA polymerase consensus sequence encodes guanosines at positions +1, +2, and +3 from the 5’ cap (40), which would severely restrict the variability of the juxtacap nucleotides in the library of motifs (Supplementary Figure 1A). We cloned the Rps20 5’ UTR downstream of a modified T7 promoter without guanosines at positions +1, +2, and +3, but were unable to synthesize mRNA from this template (data not shown). Instead, we constructed a template encoding only a single guanosine at position +1 (Supplementary Figure 1A). We introduced random nucleotides at positions +2 through +6 by PCR of the entire 5’ UTR downstream of the modified T7 promoter, and transcribed the randomized 5’UTR *in vitro* (Figure 1A). We capped the 5’UTRs *in vitro*, and performed a series of enzymatic steps to remove any uncapped RNA (Figure 1A). The library contains 1024 motifs of five randomized nucleotides downstream of the ^me7^G cap and the +1 guanosine, and prepended to the remainder of the murine Rps20 5’ UTR.

We incubated the input library with increasing concentrations of recombinant streptavidin binding peptide (SBP)-tagged eIF4E, and isolated eIF4E-bound RNAs through capture of eIF4E with streptavidin-coated magnetic beads. We constructed sequencing libraries from the bound RNAs at each concentration of eIF4E and from the input library, and performed high-throughput sequencing. We normalized the frequency of each RNA species bound to eIF4E by its frequency in the input library to identify enriched and depleted motifs.

**Figure 1:**
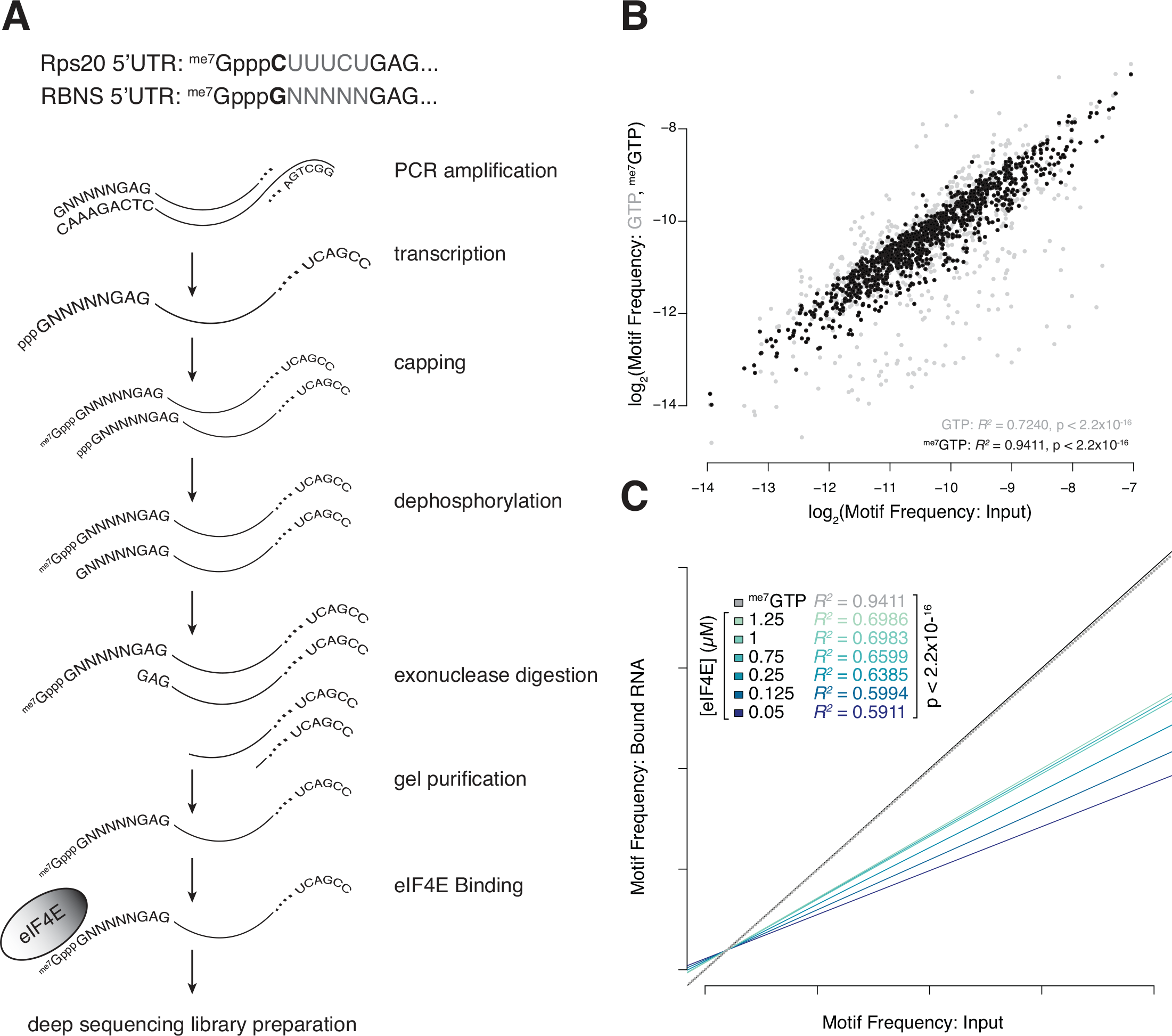
Experimental design for eIF4E Bind-n-Seq. A. Overview of RNA Bind-n-Seq library generation. Modifications to the Rps20 5’UTR and processing steps for generation of the RNA Bind-n-Seq library are shown. B. eIF4E possesses cap-dependent juxtacap nucleotide preferences. The log_2_-transformed frequency of each juxtacap motif in the input library was plotted against either its frequency in the control sample containing 1.2 5μM eIF4E and either 10 mM GTP (gray) or^me7^ GTP (black). Pearson correlation coefficients and p-values are indicated. C. Juxtacap nucleotide preferences are more apparent when eIF4E is more limiting. Linear regression was performed for the frequency of each motif in the input library versus its frequency in the eIF4E-bound library at each concentration of eIF4E, and the best-fit lines were plotted. Pearson correlation coefficients and p-values are indicated.

Although the interaction of most RNAs with eIF4E requires the 5’ cap, some RNA sequences can bind eIF4E in a cap-independent manner (41,42). To control for cap-independent and background binding, we measured binding at the highest eIF4E concentration (1.25 μM) in the presence of 7-methyl-GTP, (^me7^GTP), a competitive inhibitor of eIF4E binding to the 5’ cap, or GTP, which does not compete with the cap for binding to eIF4E. As expected, ^me7^GTP addition blocked most RNA binding to eIF4E (data not shown), suggesting predominantly cap-dependent binding. Furthermore, the motif frequencies in the ^me7^GTP-treated sample correlated very highly with input frequencies (*R^2^* = 0.9411), suggesting that the low amount of background binding we observed is sequence-independent and instead depends primarily on the frequency of each motif in the input library (Figure 1B). In contrast, we observed less correlation between the frequency of motifs in the eIF4E-bound sample and the input library for the sample containing 1.25 μM eIF4E and GTP to control for the presence of free nucleotide (*R^2^* = 0.7420) (Figure 1B), suggesting that eIF4E does have varying affinity for different motifs. In the experimental set of samples containing various concentration of eIF4E in the absence of GTP or ^me7^GTP, motif representation at the lowest eIF4E concentration (50 nM) was poorly correlated with representation in the input library (*R^2^* = 0.5911). The correlation increased with increasing eIF4E concentration (Figure 1C), which is expected as eIF4E becomes less limiting and there is less competition between RNAs for binding to eIF4E. Together, these data indicate that eIF4E does indeed bind particular juxtacap nucleotide sequences differently.

### Bind-n-Seq Experiment Reveals Juxtacap Sequence Preferences for eIF4E

In order to compare eIF4E binding between motifs present at different initial frequencies in the input library, we calculated an enrichment score. We defined the enrichment score as the frequency of each eIF4E-bound motif, relative to the frequency of the same motif in the input library. We calculated the enrichment score at all concentrations of eIF4E for the 1024 motifs present in the input library (Supplementary Table 1). To quantify the total amount of binding across all eIF4E concentrations, we calculated the area under the curve when we plotted eIF4E concentration versus enrichment score (Figure 2A). The motif with the greatest cumulative binding to eIF4E had a calculated area under the curve that was approximately 78-fold higher than the motif with the least binding to eIF4E, corresponding to a 67-fold higher enrichment score at the lowest eIF4E concentration (Supplementary Figure 2A). We performed kmeans clustering of the enrichment score at 50 nM eIF4E and examined the juxtacap sequence motifs assigned to each of the ten clusters (Supplementary Figure 2B and Supplementary Table 1). From these clusters, we classified two groups of motifs: “low binders,” which showed very little eIF4E binding, even at high eIF4E concentrations, and comprise the lowest enrichment kmeans cluster (n=77; median enrichment score for 50 nM eIF4E = 0.308, median AUC = 1.27), and “high binders,” which exhibited high eIF4E binding, even at low eIF4E concentrations, and comprise the highest three enrichment kmeans clusters (n = 46; median enrichment score for 50 nM eIF4E = 3.23; median AUC = 14.34) (Figure 2B and 2C). Motifs with poor eIF4E binding were highly enriched for guanosine in the +2 position and depleted for guanosine in the +6 position, with no other major nucleotide preferences (Figure 2C and Supplementary Figure 2C and 2E). In contrast, enrichment of +4 guanosine and +5 cytosine, and depletion of +6 adenosine were characteristic of the high binding motifs (Figure 2C and Supplementary Figure 2D and 2E). These data indicate that eIF4E has position-dependent preferences for juxtacap nucleotides.

**Figure 2:**
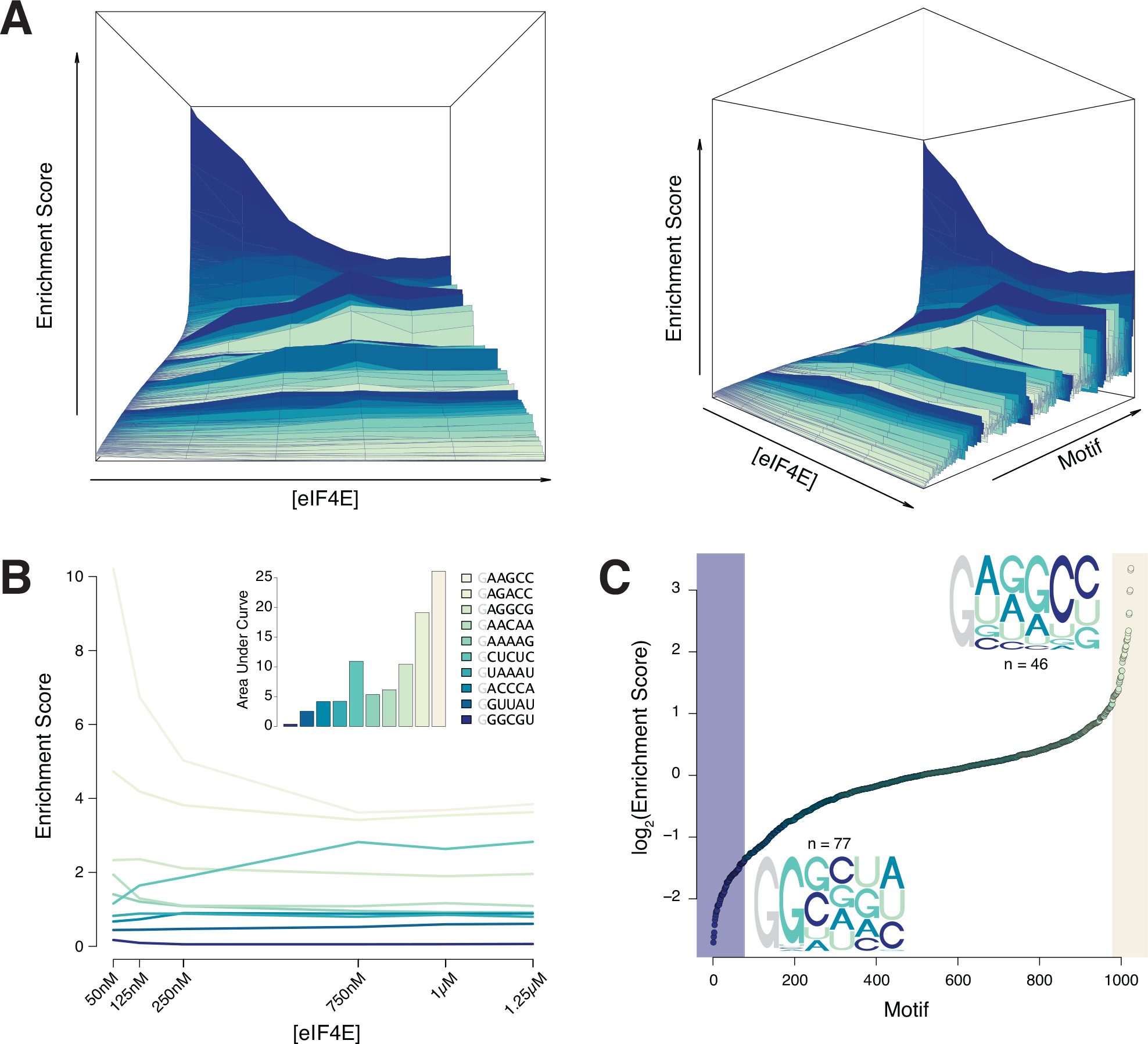
eIF4E Bind-n-Seq reveals specific juxtacap sequence preferences. A. Overview of the distribution of enrichment scores for each motif across all eIF4E concentrations. Motifs were sorted from least enriched (front) to most enriched (back) in the 50nM eIF4E sample, and each eIF4E concentration was plotted against the enrichment score at that eIF4E concentration for each motif. B. Examples of enrichment score distributions and area under the curve for individual motifs from each kmeans group across all eIF4E concentrations. Main: eIF4E concentration is plotted against enrichment score for individual motifs from each kmeans group. Inset: Area under the curve was plotted for each representative motif. C. Distribution of enrichment scores at 50nM eIF4E and nucleotide frequencies for high-and low-eIF4E-binding motifs reveals nucleotide preferences. The log_2_ of the enrichment score at 50nM eIF4E was plotted for each motif. Highlighted are the lowest kmeans group (n=77) and highest three kmeans groups (n=46). Juxtacap motifs for the lowest and three highest kmeans groups are indicated.

### The Identity of the Juxtacap Nucelotides Affects Translation

Having established that juxtacap sequence identity influences eIF4E binding, we sought to determine whether the juxtacap sequence also influences translation. We individually cloned 5’UTRs of the Rps20 gene containing juxtacap motifs from the clusters of low and high eIF4E binding motifs downstream of the modified T7 promoter and upstream of a *Renilla* luciferase-expressing construct harboring an encoded polyA tail (Figure 3A). These constructs contain a 5’ UTR that is identical to those in the library used for RNA Bind-n-Seq except for the Kozak sequence, but encode *Renilla* luciferase so that we could measure the level of translation of each mRNA in a cell-free translation assay. We *in vitro* transcribed and cotranscriptionally capped the RNAs using anti-reverse cap analog, and measured their translation *in vitro* in HeLa cell lysates. Indeed, low eIF4E binders were translated more poorly than high eIF4E binders (Figure 3B), suggesting that the juxtacap sequence identity can modulate translation.

**Figure 3:**
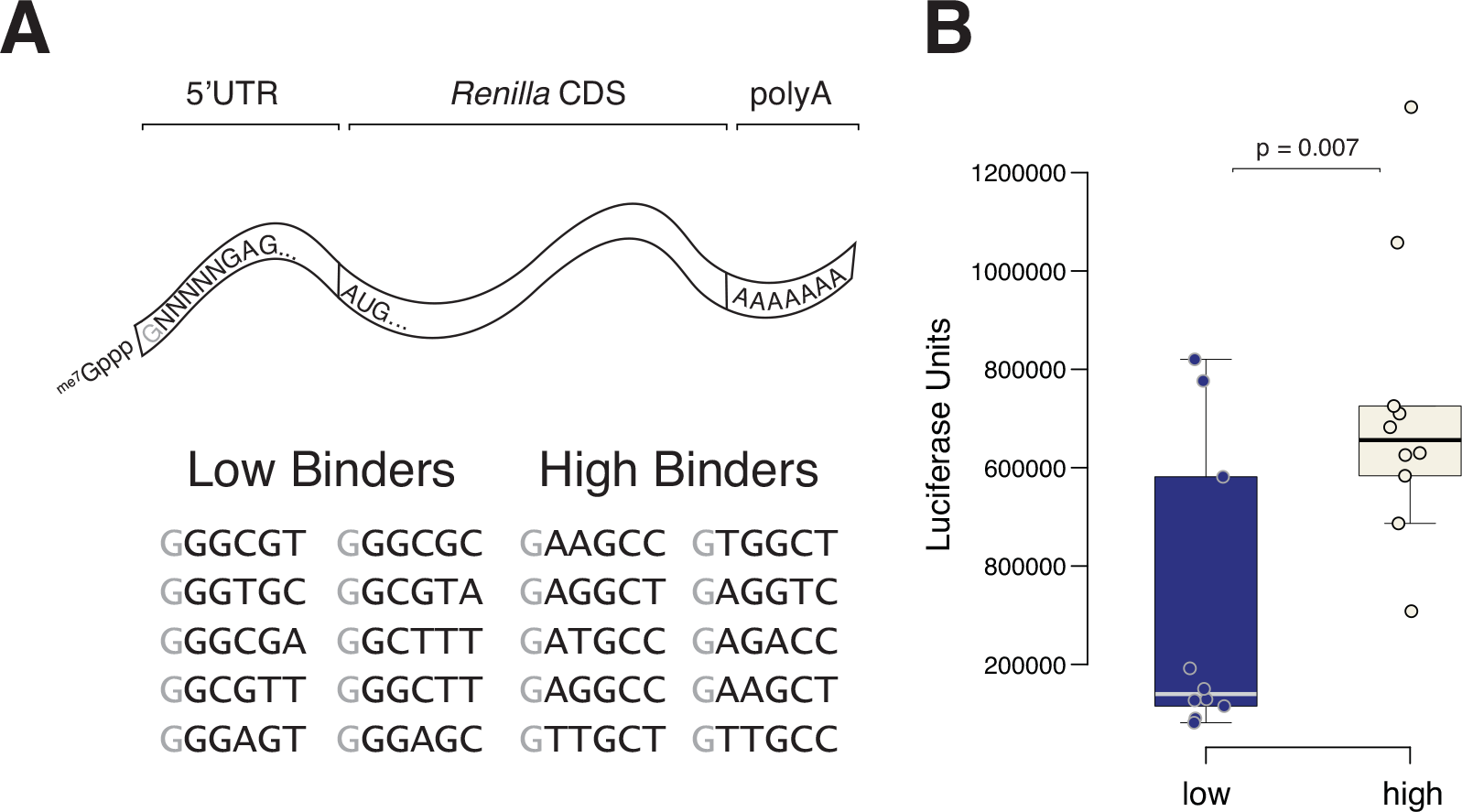
Juxtacap nucleotide identity affects translation. A. Diagram of the *Renilla* luciferase-expressing construct and list of low-and high-eIF4E-binding motifs whose translation was measured. B. Low-and high-binding motifs are translated differently in a cell-free translation system. Box-and-whisker plots for low-eIF4E-binding and high-eIF4E-binding motifs were generated from the mean luciferase unit values from three independent experiments for each mRNA. The mean luciferase values for each mRNA are indicated as points. Significance was determined by the Wilcoxon rank sum test.

### Juxtacap Nucleotide Identity Does not Correlate with Translation in Cells

The observation that eIF4E possesses a range of binding preferences for RNAs containing different juxtacap sequences suggests two alternative hypotheses. The first possibility is that eIF4E directly interacts with nucleotides downstream of the 5’ cap, and that the juxtacap nucleotide sequence directly dictates the extent of eIF4E interaction; in this case, the effect of a juxtacap nucleotide sequence would be maintained across all mRNAs containing that juxtacap nucleotide sequence and translated in a cap-dependent fashion, regardless of the 5’UTR context. An alternative possibility is that eIF4E does not directly recognize the nucleotides downstream of the cap, but rather is affected by RNA secondary structures adjacent to the cap, which exist in the context of the longer 5’UTR and remainder of the mRNA. If this were the case, juxtacap motifs would promote specific RNA structures, which would dictate the level of eIF4E binding and subsequent translation. These structures would be independent of the specific sequence of the juxtacap nucleotides, and instead would rely on the greater sequence context of the juxtacap motifs.

To investigate the first possibility, we compared published transcriptional start site (36) and ribosome footprinting (21) datasets and asked whether our low-and high-eIF4E-binding motifs were associated with lowly-and highly-translated mRNAs in cells, respectively. Importantly, we examined only endogenous 5’UTRs that begin with a guanosine, as this feature was invariant in our RNA Bind-n-Seq library. We observed no overall correlation between ribosome occupancy and binding (Figure 4A), and no difference in the ribosome occupancy across kmeans enrichment score groups (Supplementary Figure 3A). Furthermore, we saw no difference when we compared the high-binding motifs represented in the ribosome footprinting dataset (top three kmeans enrichment score clusters; 42/46 motifs mapped to 210 endogenous mRNAs) with the low-binding motifs represented (lowest kmeans enrichment score group; 53/77 motifs mapped to 2763 endogenous mRNAs) (Figure 4B), nor when we compared both low-and high-binding groups to the rest of the mapped motifs (Figure 4B). Additionally, we performed kmeans clustering by ribosome occupancy (Figure 4C, Supplementary Figure 3B, and Supplementary Table 2), and examined the juxtacap nucleotide preferences of highly-and lowly-translated motifs. Neither group of motifs showed significant nucleotide preferences (Figure 4C), further indicating that the effect of juxtacap sequence on translation is context-dependent. These analyses suggest that the juxtacap sequence itself is not sufficient to restrict or promote mRNA translation in cells.

**Figure 4:**
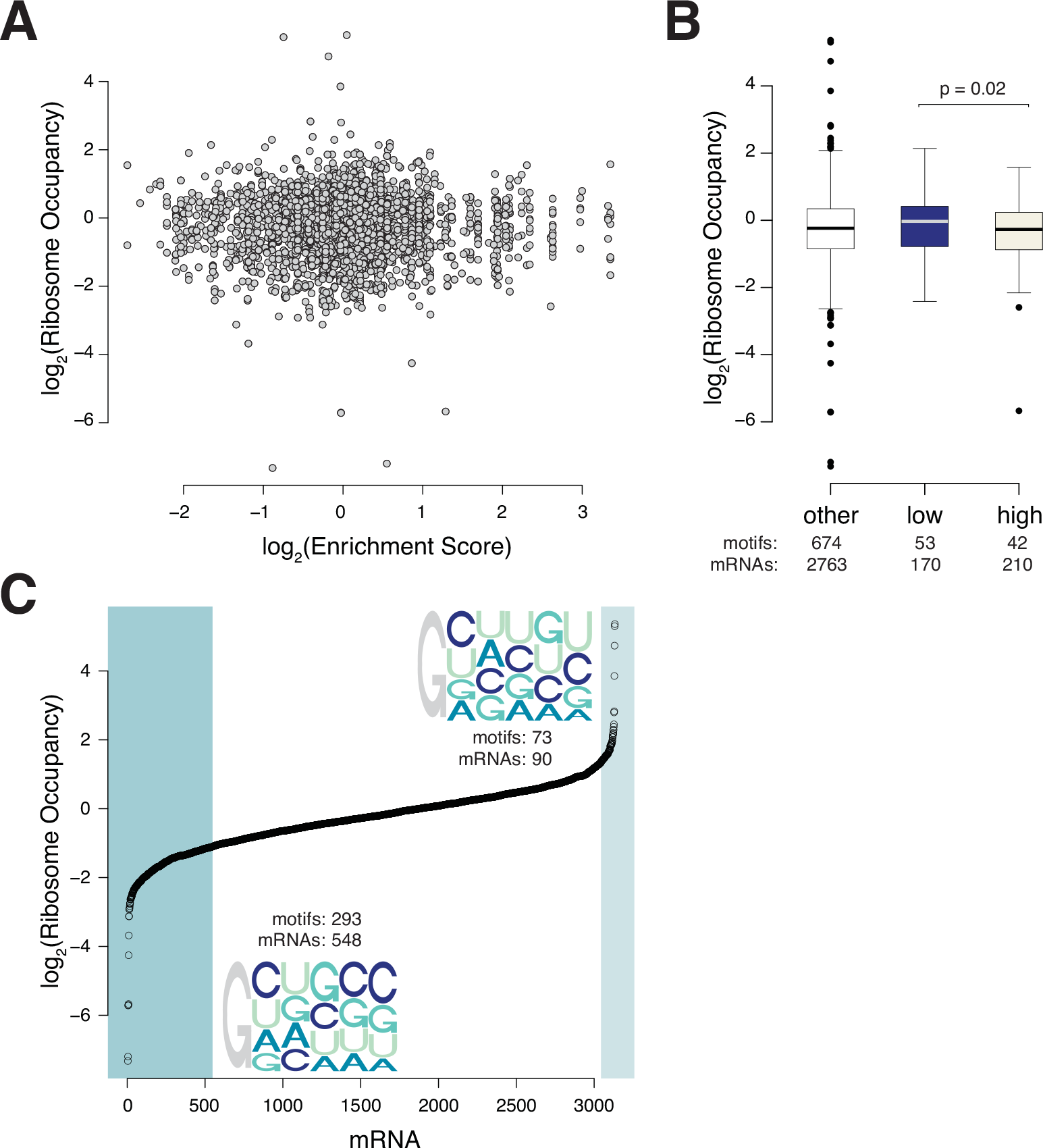
Juxtacap nucleotide identity does not correlate with mRNA translation in cells. A. Ribosome occupancy of endogenous mRNAs that begin with a +1 guanosine does not correlate with eIF4E binding. The log_2_ Bind-n-Seq enrichment score for each motif at 50nM eIF4E is plotted against the log_2_ ribosome occupancy of endogenous mRNAs. B. Ribosome occupancy of endogenous mRNAs containing low-and high-eIF4E-binding juxtacap motifs is not different from ribosome occupancy of mRNAs containing other juxtacap motifs. Log_2_ ribosome occupancy of endogenous mRNAs containing low-binding, high-binding, or other motifs is plotted. C. Highly-and lowly-translated endogenous mRNAs do not exhibit nucleotide preferences. The distribution of log_2_ ribosome occupancy was plotted. Highlighted are the lowest kmeans group (n = 293 motifs represented in 548 mRNAs) and four highest kmeans groups (n = 73 motifs represented in 90 mRNAs). The juxtacap motifs for lowest and four highest kmeans groups are indicated.

### Juxtacap Nucleotide Sequences Can Modulate Translation by Defining the Cap-Proximal RNA Structure

The lack of correlation between the motifs enriched in eIF4E binding from our Bind-n-Seq and translation of mRNAs in cells suggests that the effect of cap-proximal nucleotide identity on eIF4E binding is mRNA-context-dependent. We sought to test the notion that juxtacap nucleotide identity can define a particular structure that restricts or promotes translation in a constrained 5’UTR context. Secondary structure prediction (39) for the high and low eIF4E-binding groups in the context of the Rps20 5’UTR revealed striking differences in the accessibility of the 5’ end of the 5’UTRs containing high-and low-eIF4E-binding motifs. Specifically, low-binding motifs were predicted to have more highly base paired cap-proximal nucleotides, while high-binding motifs had more open juxtacap sequences (Figure 5A, Supplementary Figure 4A and 4B). The base pairing interactions in both the high-and low-eIF4E-binding groups were dependent on nucleotide identity downstream of the juxtacap sequence.

**Figure 5:**
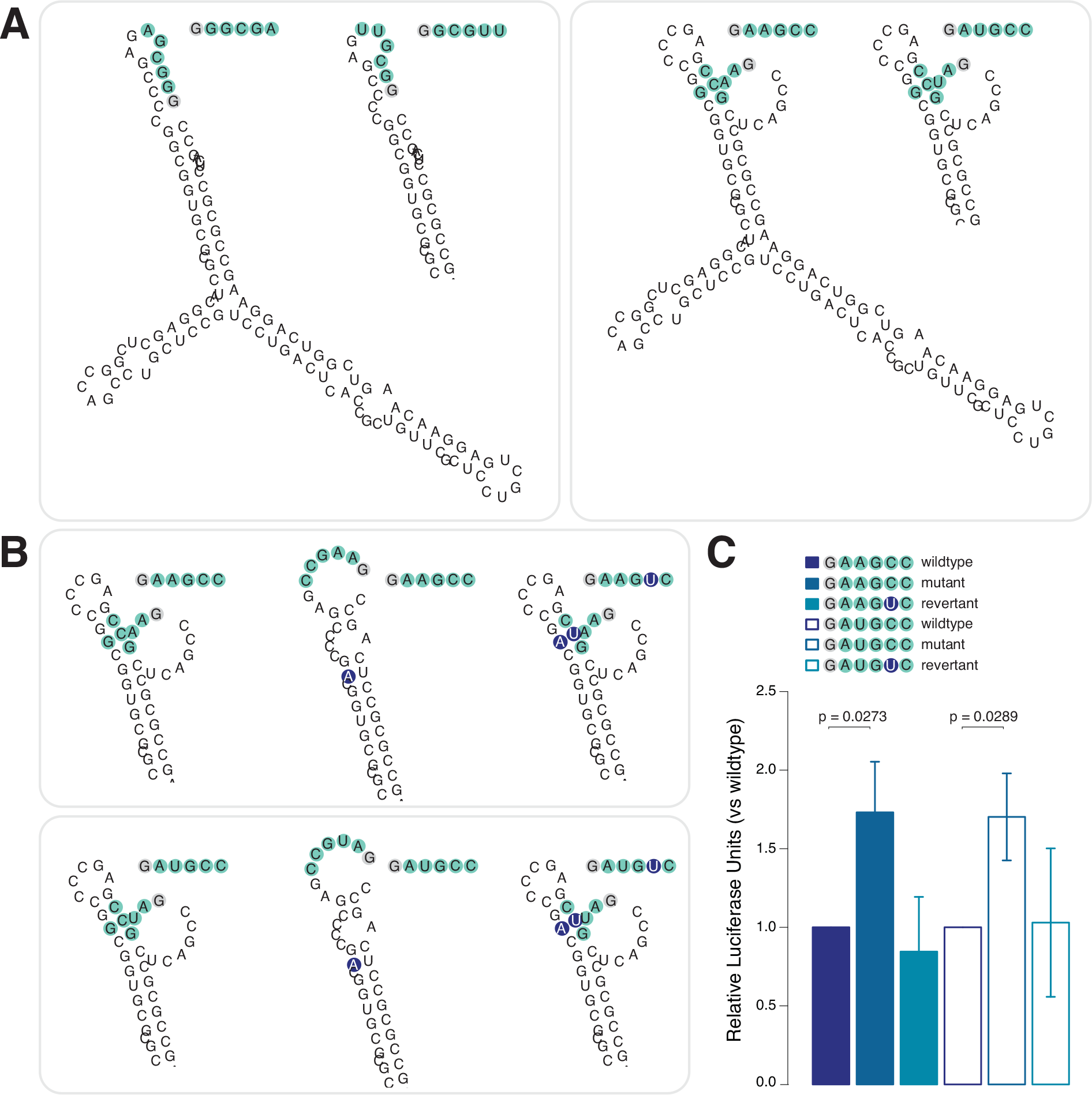
Juxtacap nucleotide sequence modulates translation by defining cap-proximal structure. A. The predicted minimum free energy (MFE) structures of mRNAs containing low-eIF4E-binding motifs have less accessible 5’ ends than those contatining high-eIF4E-binding motifs. MFE structures for mRNAs with two low-binding motifs and two high-binding motifs are shown. The portion of the structure not shown for the second mRNA in each group is identical to that region in the first mRNA. The juxtacap motif nucleotides are highlighted. B. Mutations downstream of the juxtacap sequence are predicted to make the 5’ end of mRNAs containing high-eIF4E-binding motifs more accessible. The 5’ end of the predicted MFE structures for mRNAs with wildtype, mutant, and revertant juxtacap motifs for two high-eIF4E-binding motifs are shown; the portion of the structures not shown for each mRNA are identical to that region in the wildtype mRNA shown in Figure 5A. The juxtacap motif nucleotides are highlighted, with mutated nucleotides indicated in purple. C. Mutations downstream of the juxtacap sequence predicted to make the 5’ end of mRNAs containing high-eIF4E-binding motifs more accessible also increase translation. The mutant and revertant luciferase unit values for each replicate were normalized to the corresponding wildtype values, and mean +/-SEM of the normalized value for three independent experiments were plotted. Significance was determined by a one-tailed Student’s t-test with equal variances on non-normalized values (variances were determined to be equal using the F test). P-values are indicated.

In the low-binding group, the +2 guanosine characteristic of the low-binding motifs is predicted to be base paired (Figure 5A and Supplementary Figure 4A) and is associated with concurrent base pairing at the invariant +1 guanosine (Figure 5A). This base pairing could severely restrict eIF4E binding to the cap and subsequent translation. In contrast, the high-binding motifs were predicted to contain several free juxtacap nucleotides, and the nucleotide with the strongest position-dependent preference, +5 cytosine, is predicted to participate in a base pairing interaction with a downstream guanosine at position +15 of the 5’ UTR (Figure 5B and Supplementary Figure 4B).

We mutated this downstream guanosine to an adenosine to disrupt the base pairing interaction and increase the accessibility of the juxtacap sequence (Figure 5B), and measured translation in a cell-free translation system (Figure 5C). Mutation of the downstream guanosine in two high-eIF4E-binding Rps20 5’UTRs containing motifs predicted to form a base pair between the +5 cytosine and +15 guanosine increased translation, while the compensatory mutation at position +5 in the juxtacap motif decreased translation (Figure 5B and 5C). Modifying the Watson-Crick base pairing between the Rps20 5’UTR and the juxtacap nucleotides can alter translation, suggesting that the juxtacap sequence influences eIF4E binding and translation by defining the accessibility of the 5’ end of the mRNA.

## DISCUSSION

Initiation is normally the rate-limiting step of translation, and is tightly regulated via multiple mechanisms (reviewed in (4)). Molecularly defining the preferences of initiation factors for particular mRNAs is important not only for predicting how efficiently a given mRNA will be translated, but also for understanding how different regulatory inputs restrict or promote the translation of specific messages. There is an abundance of structural and biophysical evidence describing how eIF4E binds analogs of the 5’ cap (22-27). However, there was previously very little understanding of how juxtacap nucleotides affect eIF4E binding and subsequent translation.

Studies that thoroughly characterize the 5’ ends of mRNA transcripts have uncovered that, for many genes, transcription does not begin at a single well-defined position, but can often occur at multiple discrete sites, or as a distribution around a particular site (43,44). In some cases, these alternative transcriptional start sites (TSSs) substantially affect the 5’UTR of the mRNA, which can have considerable consequences for translation (45). However, it has remained unknown whether the difference of a few nucleotides adjacent to the 5’ cap can influence translation. Our findings suggest that the juxtacap sequence can influence the accessibility of the 5’ end of an mRNA, and that small changes in this sequence can modulate translation. Our model implies that transcripts encoded by genes possessing even a narrow distribution of TSSs may be translated quite differently from one another. In the case of a few genes with multiple discrete TSSs, selection of a particular TSS is regulated by transcription factor expression (43,46). Although it is unlikely that TSS selection for genes possessing a distribution of TSSs around a particular site would rely on different transcription factors, it is an attractive notion that these distributed TSSs could also be regulated. Regardless of their regulation, we predict that even small differences in TSSs can influence eIF4E binding and translation.

In cells, eIF4E is limiting (13,14), and therefore is bound to either eIF4G or a 4EBP. Binding of an eIF4G fragment that associates with eIF4E but does not contain the RNA binding domains of eIF4G can alter the structure of eIF4E and increase its affinity for the 5’ cap (47,48). Although so far this RNA-binding-independent conformational coupling has only been directly observed in yeast (24,47,48), it raises the possibility that the juxtacap sequence preferences of eIF4E when bound to eIF4G or a 4EBP could be different than for eIF4E alone. We also predict that some differences in eIF4E binding dictated by the juxtacap sequence are masked *in vivo*, due to stabilization of the eIF4E-mRNA interaction by direct binding of eIF4G to the RNA (49,50). Furthermore, juxtacap motifs that reduce eIF4E binding primarily through increasing the off-rate of eIF4E would be particularly sensitive to the presence of eIF4G, because eIF4G could act to keep eIF4E tethered to the mRNA after initial binding. The lowly-and highly-bound motifs we identified by RNA Bind-n-Seq were translated differentially, suggesting that the presence of eIF4G does not fully conceal all eIF4E-dependent juxtacap binding differences. However, the possibility remains that particular juxtacap sequences could influence eIF4E binding differently in the presence of eIF4G.

The nucleotide adjacent to the cap, corresponding to the first transcribed nucleotide, can directly contact eIF4E (25), and reduces the affinity of the 5’ cap for eIF4E in biochemical assays (26,51). In mammals, the ribose of the +1 nucleotide is typically methylated at the 2’ position, and often the +2 nucleotide is methylated as well (52,53). 2’-*O*-methylation was first identified in tRNA and in functional regions of rRNA (reviewed in (54,55)), and this modification is thought to alter RNA structure, RNA-RNA interactions, and RNA-protein interactions (reviewed in (56)). Although the functions of this modification have yet to be elucidated for most mRNAs, it may play a role in distinguishing self from non-self RNA (57,58). While knockdown of the enzyme responsible for 2’-*O*-methylation at the +1 position did not affect global translation in HeLa cells (59), it was shown for specific mRNAs that 2’-*O*-methylation of the first two nucleotides can affect ribosome binding and translational efficiency (60-63). It is probable that 2’-*O*-methylation alters translational efficiency at least in part via effects on eIF4E binding, given that at least the modified +1 nucleotide can contact eIF4E directly. It is also possible that modification of these nucleotides could alter binding of eIF4E differently in the context of different juxtacap sequences, which would allow mRNAs to display even greater variation in eIF4E recruitment and translation than by differences in the juxtacap sequence alone.

The eIF4E used in our study was prepared from *E. coli*, and hence was unphosphorylated. In mammals, eIF4E is phosphorylated by two kinases that bind eIF4G, Mnk1 and Mnk2 (64,65). In the presence of ample nutrients and growth factors, mTORC1 is active and phosphorylates the 4EBPs (10,11). This event liberates eIF4E and allows it to bind eIF4G, and subsequently become phosphorylated by Mnk1/2 at serine 209. The phosphorylation site is located within the flexible C-terminal loop of eIF4E that is stabilized by the presence of a +1 adenosine in the cocrystal structure (25). It is positioned directly adjacent to the cap-binding pocket of eIF4E, a prime location for influencing juxtacap nucleotide recognition. Although the precise molecular function of this phosphorylation is not thoroughly understood ((30,51,66,67); reviewed in (68)), this phosphorylation site is clearly important for translation of a subset of mRNAs that promote transformation and tumorigenesis (69). These mRNAs are distinct from the mRNAs whose translation is promoted by wildtype eIF4E overexpression (19), and it is likely that phosphorylated eIF4E and unphosphorylated eIF4E possess different juxtacap nucleotide preferences.

Although the effect of juxtacap sequence on eIF4E binding and translation was primarily mRNA context-dependent in our study, we do not rule out the notion that eIF4E can directly recognize the juxtacap nucleotides. Importantly, our experimental constraints do not allow us to assess the effect of differences in the first juxtacap nucleotide, while structural and biophysical evidence suggests nucleotides at this position are of particular interest (22,25,26,51). Nucleotides in the +1 position can form different contacts with eIF4E and alter the affinity of eIF4E for the 5’ cap, depending on the nucleotide identity (25,51), and we predict that differences in the +1 nucleotide will alter the juxtacap sequence preferences of eIF4E. To fully assess the role of the juxtacap sequence in eIF4E binding and translation, it will be important to develop methods to readily synthesize capped mRNAs encoding different +1 nucleotides; for example, by improving existing chemical synthesis methods (70) or by identifying an RNA polymerase that can produce mRNAs with various +1 nucleotides and is adaptable to robust *in vitro* synthesis. A third possibility would be to identify enzymes that can phosphorylate RNA 5’ ends to produce 5’-triphosphorylated RNA (71), which could be used as a substrate for existing *in vitro* capping systems.

In addition to its importance for cap-dependent translation, eIF4E is associated with several other processes in the cell. For instance, eIF4E is involved in the export of particular mRNAs from the nucleus (reviewed in (72)). eIF4E is also a component of stress granules and P bodies (reviewed in (73)), which serve as mRNA storage compartments under conditions of stress and as sites of mRNA degradation, respectively. Cap-proximal nucleotide preferences could potentially contribute to mRNA selection for any of these other processes involving eIF4E.

The idea that mRNA affinity for the translational machinery can modulate translation is not new, although there are few specific examples of this phenomenon. Over forty years ago, Lodish proposed the model that translation of mRNAs that initiate protein synthesis at lower rates will be preferentially inhibited when initiation is globally reduced, which was true when comparing translation of alpha and beta globin (74). Almost a decade later, a difference in the affinity for the general translation initiation factor eIF2 was experimentally shown to mediate selective translation of a particular viral mRNA over globin mRNA in a cell-free system (75). Decades of work on mRNAs containing internal ribosome entry sites (IRESs) has shown that they rely on direct recruitment of general translation initiation factors or even the ribosome for their translation (reviewed in (76,77)). Here, we describe the direction, magnitude, and sequence-dependence of the effect of the juxtacap nucleotides on eIF4E binding to a model 5’UTR. We show that juxtacap sequence identity can affect translation, likely by contextually defining cap-proximal end accessibility. Our work increases the understanding of how mRNAs are chosen for translation, and raises the possibility that initiation factor preferences are a more widespread mechanism for dictating translational efficiency of an mRNA than previously appreciated.

## SUPPLEMENTARY DATA

Supplementary data are available at NAR Online.

## ACKNOWLEDGEMENTS

We would like to thank Larry Schweitzer for critical reading of the manuscript and Carson Thoreen for generous sharing of custom analysis scripts, as well as invaluable input from L.S. and C.T. throughout all stages of this work. We would like to thank Ellen Edenberg for critical reading of the manuscript, Bingbing Yuan for CAGEr analysis, and the entire Sabatini Lab for helpful feedback. We would also like to thank the Whitehead Institute Genome Technology Center for sequencing.

